# Optimization of T-DNA architecture for Cas9-mediated mutagenesis in Arabidopsis

**DOI:** 10.1101/419952

**Authors:** Baptiste Castel, Laurence Tomlinson, Federica Locci, Ying Yang, Jonathan D G Jones

## Abstract

Bacterial CRISPR systems have been widely adopted to create operator-specified site-specific nucleases. Such nuclease action commonly results in loss-of-function alleles, facilitating functional analysis of genes and gene families We conducted a systematic comparison of components and T-DNA architectures for CRISPR-mediated gene editing in Arabidopsis, testing multiple promoters, terminators, sgRNA backbones and *Cas9* alleles. We identified a T-DNA architecture that usually results in stable (*i.e.* homozygous) mutations in the first generation after transformation. Notably, the transcription of sgRNA and Cas9 in head-to-head divergent orientation usually resulted in highly active lines. Our Arabidopsis data may prove useful for optimization of CRISPR methods in other plants.

## INTRODUCTION

CRISPR (clustered regularly interspaced short palindromic repeat)-Cas (CRISPR associated) site-specific nucleases evolved as components of prokaryotic immunity against viruses, and are widely deployed as tools to impose operator-specified nucleotide sequence changes in genomes of interest [1–4]. During infection by bacteriophages, Cas1 and Cas2 can integrate phage DNA sequences into ‘spacer’ regions of tandem CRISPR loci in the bacterial genome. The crRNA (CRISPR-RNA) transcription product of the spacer associates with nucleases from the Cas family to form ribonucleoproteins that can cleave nucleic acid sequences homologous to the spacer. This enables elimination of viral nucleic acid upon subsequent infection. CRISPR systems are divided in two classes [5,6]. Class 1 systems comprise multi-subunit complexes whereas Class 2 systems function with single ribonucleoproteins. Within Class 2, Type-II and Type-V cleave dsDNA (double-stranded DNA) via Cas9 and Cas12/Cpf1 respectively, while Type-VI cleaves ssRNA (single-stranded RNA) via Cas13/C2c2.

Cas9, Cas12 and Cas13-based systems function in heterologous organisms enabling applications such as targeted mutagenesis, dynamic imaging of genomic loci, transcriptional regulation, pathogen detection and RNA quantification [7–9]. Expression of Cas9 with its associated sgRNA (single-guide RNA, an artificial fusion of the dual endogenous crisprRNA/trans-acting-crisprRNA), results in targeted DNA mutations in animals and plants [3,10,11]. Cas9-sgRNA ribonucleoprotein cleaves genomic DNA at loci homologous to the sgRNA spacer sequence. Cleaved DNA strands can be religated by the endogenous Non-Homologous End Joining (NHEJ) system, which can result in insertions or deletions (indels) at the repaired site. Indels in the CDS (coding DNA sequence) can cause a codon reading frame shift resulting in loss-of-function alleles.

*Arabidopsis thaliana* (Arabidopsis) is widely used for plant molecular genetics. Expression of CRISPR-Cas9 components can result in loss-of-function alleles of targeted genes in Arabidopsis, with variable efficiency [12–14]. To improve induced mutation rates in Arabidopsis, several groups have evaluated various promoters to drive *Cas9* expression. [15–17].

We set out to optimize mutation rates in Arabidopsis, and report here an extensive comparison of promoters, *Cas9* alleles, terminator, sgRNA and construct architecture. Cas9-sgRNA ribonucleoprotein can be directly delivered by protoplast transformation or particle bombardment into plant cells [18,19], but these methods require regeneration via tissue culture. To avoid this process, we delivered *Cas9* and the *sgRNA* in transgenic Arabidopsis. This method requires three steps: (i) DNA assembly of a binary vector with selectable marker, a *Cas9* and a *sgRNA* expression cassettes in the T-DNA, (ii) *Agrobacterium tumefaciens*-mediated stable transformation of the plasmid via the floral dip method [20] and (iii) identification of mutants among the transformed lines. Multiple T-DNA architectures were tested for their ability to trigger homozygous mutations in the *ADH1* gene, including presence or absence of an “*overdrive*” sequence to promote T-DNA transfer [21]. ADH1 converts allyl alcohol into lethal allyl aldehyde, so *adh1* mutant lines resist allyl-alcohol treatment, enabling facile measurement of CRISPR-induced mutation rates [13,16]. We defined combinations of CRISPR components that enable high efficiency recovery of stable homozygous mutants in one generation.

## RESULTS

### Golden Gate cloning enables facile assembly of diverse Cas9 T-DNA architectures

In Golden Gate modular cloning, the promoter, reading frame and 3’ end modules at ‘Level 0’, are assembled using Type IIS restriction enzymes to ‘Level 1’ complete genes, that can then be easily combined into T-DNAs carrying multiple genes at ‘Level 2’. This enables facile assembly of diverse T-DNA conformations [22,23]. Level 0 acceptor vectors are designed to clone promoter, coding sequence (CDS) or terminator fragments (see Materials and Methods). For our purpose, we used three Level 1 vectors: a glufosinate plant selectable marker in position 1 (pICSL11017, cloned into pICH47732), a *Cas9* expression cassette in position 2 (cloned into pICH47742) and a *sgRNA* expression cassette in position 3 (cloned into pICH47751) (Fig 1). Some *Cas9* expression cassettes were cloned into a Level 1 position 2 variant: pICH47811. This vector can be assembled in Level 2 in the same fashion as pICH47742, but it enables *Cas9* transcription in the opposite direction as compared to the other Level 1 modules. We assembled 25 different Level 1 *Cas9* constructs and four *sgRNA* expression cassettes. The sequence targeted by the sgRNA was CGTATCTTCGGCCATGAAGC(*NGG*) (Protospacer Adjacent Motif indicated in italics) which targets specifically *ADH1* in Col-0, enabling pre-selection of CRISPR-induced *adh1* mutants by selecting with allyl alcohol [13]. Assembly of these Level 1 modules resulted in 39 Level 2 T-DNA vectors (Table S1). We used two types of Level 2 vectors, with an *overdrive* (pICSL4723) or without (pAGM4723). We found that it does not affect the CRISPR-induced mutation rates (Fig S1). Thus, constructs transformed in pICSL4723 or in pAGM4723 can be compared side-by-side. More details of the assembly protocols can be found in the ‘Materials and Methods’ section.

**Fig 1:**
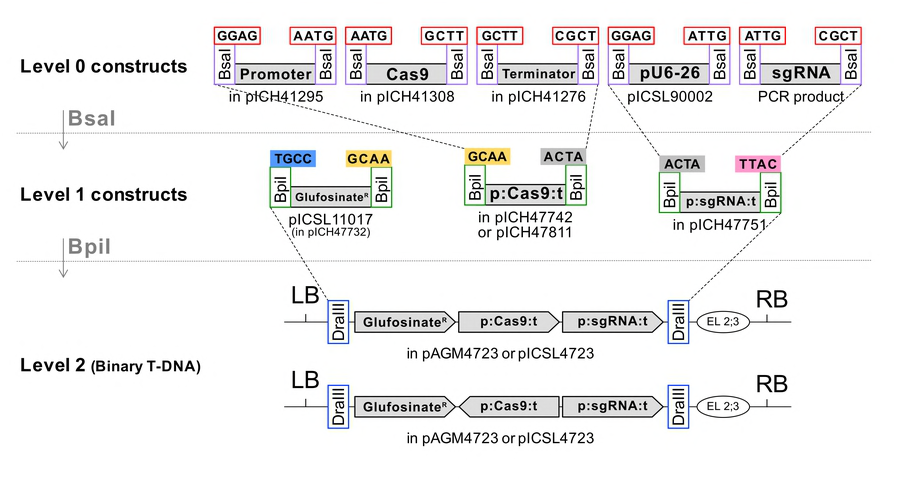
Golden Gate cloning method enables assembly of CRISPR modules in various combinations. *Cas9* alleles, promoters and terminators were cloned into the indicated Level 0 acceptor vectors as described in *Materials and Methods* and were assembled in Level 1 acceptor vector pICH47742. sgRNAs targeting *AtADH1* were amplified by PCR and assembled with the *U6-26* promoter vector pICSL90002 in the same manner. Both Cas9 and sgRNA expression units were assembled in Level 2 acceptors pAGM4723 (not containing an *overdrive* sequence) or pICSL4723 (containing an *overdrive*) along with a Glufosinate resistance plant selectable marker. An end-linker pICH41766 (EL2;3) was used to link the sgRNA expression unit to the Level 2 acceptor vector. For a “head-to-head orientation” of the sgRNA and Cas9 expression cassettes, Cas9 allele, promoter and terminator were assembled into pICH47811 instead of pICH47742.

### CRISPR-induced Arabidopsis mutations can be selected using allyl-alcohol

The 39 Level 2 plasmids were transformed in *A. tumefaciens* strain GV3101 and used to generate Arabidopsis Col-0 transgenic lines. ‘T1’ refers to independent primary transformants selected from the seeds of the dipped plant; ‘T2’ refers to the T1 progeny. For each of the 39 constructs, about 100 T2 progenies from six independent T1 lines were screened for allyl alcohol resistance (Fig 2). T2 seeds were selected with 30 mM allyl-alcohol for two hours. Six survivors (or all survivors if there were less than six) were screened by PCR amplification and capillary sequencing to confirm the mutation in *ADH1* at the expected target site. This genotyping step enabled us to estimate the percentage of non-mutated plants that escape the allyl-alcohol selection. We indeed identified some lines surviving the allyl-alcohol screen that are heterozygous (*ADH1/adh1)*. CRISPR activity is expressed as [(number of allyl-alcohol surviving plants) x (% of homozygous or biallelic mutants confirmed by sequencing among the surviving plants tested) / (number of seeds sown)]. It was measured for six independent T2 families, for each of 39 constructs. When more than 75% of the lines survived the allyl-alcohol treatment and all the lines genotyped are knock-out (KO) alleles with the exact same mutation within one T2 family, we assumed that the T1 parent was a homozygous mutant. Such T2 families are indicated in red.

**Figure 2:**
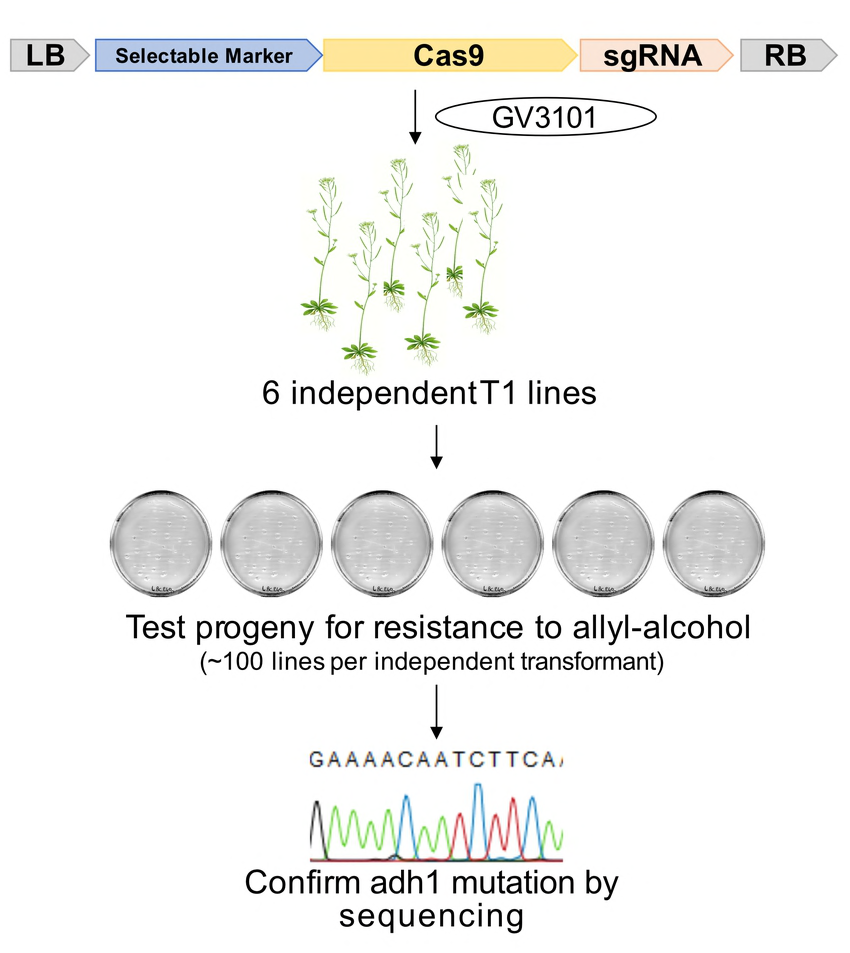
Evaluation of mutation rates. Constructs were transformed into Arabidopsis accession Col-0 via *Agrobacterium tumefaciens* strain GV3101. Six independent transformants (T1) were selected using Glufosinate. About 100 progeny (T2) of each transformant were selected for allyl-alcohol resistance. For each independent T2 family, up to six allyl-alcohol resistant plants were genotyped at the *ADH1* locus. For each T2 family, the mutation rate was calculated as [(number of allyl-alcohol surviving plants) x (% of homozygous or biallelic mutants confirmed by sequencing among the surviving plants tested) / (number of seeds sown)].

### *UBI10, YAO* and *RPS5a* promoter-controlled *Cas9* expression enhance mutation rates

CRISPR-mediated DNA sequence changes are only inherited if they occur in the germline. The Cauliflower Mosaic Virus 35S promoter and ubiquitin promoters are strongly expressed in most tissues [24]. We compared the *35S* and Arabidopsis *UBI10* promoters. More mutants were recovered using the *UBI10* promoter, suggesting it is more active than *35S* in the germline (Fig 3A). Following this observation, we tested other germline-expressed promoters.

**Figure 3:**
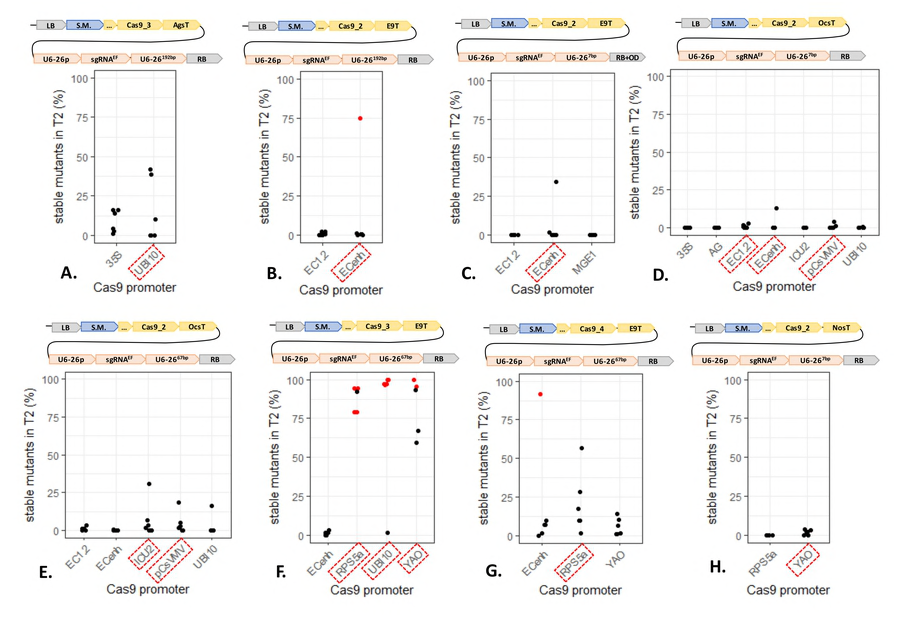
*UBI10, YAO* and *RPS5a* promoter-regulated *Cas9* expression enhances mutation rates. **A. to H.** Each panel represents a promoter comparison in the same T-DNA context. Promoters can be compared within each panel, but not from one panel to another. The modules were assembled into pICSL4723 (RB+OD, with an *overdrive*) or pAGM4723 (RB, without an *overdrive*) and transformed into Col-0 via *Agrobacterium tumefaciens* strain GV3101. <u>LB</u>: Left Border. <u>S.M</u>.: Selectable Marker (*Glufosinate resistance* gene). <u>35S</u>: 426 bp of the 35S promoter from *Cauliflower Mosaic Virus*. <u>UBI10</u>: 1327 bp of the At4g05320 promoter. <u>EC1.2</u>: 1014 bp of the At2g21740 promoter. <u>EC_enh.</u>: 752 bp of the At2g21740 promoter fused to 548 bp of the At1g76750 promoter. <u>MGE1</u>: 1554 bp of the At5g55200 promoter. <u>AG</u>: 3101 bp of the At4g18960 promoter. <u>ICU2</u>: 625 bp of the At5g67100 promoter. <u>pCsVMV</u>: 517 bp of a promoter from *Cassava Vein Mosaic* virus. <u>RPS5a</u>: 1688 bp of the At3g11940 promoter. <u>YAO</u>: 596 bp of the At4g05410 promoter. <u>Cas9_1</u>: Mali *et al.*, 2013 [3]. <u>Cas9_2</u>: Fauser *et al.*, 2014 [13]. <u>Cas9_3</u>: Li *et al.*, 2013 [25]. <u>Cas9_4</u>: Le Cong *et al.*, 2013 [10]. <u>E9T</u>: 631 bp of the *Pisum sativum rbcS E9* terminator. <u>OcsT</u>: 714 bp of the *Agrobacterium tumefaciens octopine synthase* terminator. <u>AgsT</u>: 410 bp of the *Agrobacterium tumefaciens agropine synthase* terminator. <u>NosT</u>: 267 bp of the *Agrobacterium tumefaciens nopaline synthase* terminator. <u>U6-26p</u>: 205 bp of the At3g13855 promoter. sgRNA^EF^: “extension-flip” sgRNA. <u>U6-26t</u>: 7, 67 or 192 bp of the At3g13855 terminator. <u>RB</u>: Right Border. The sgRNA targets *ADH1*. CRISPR activity measured in % of homozygous or biallelic stable mutants in the second generation after transformation (T2). Each dot represents an independent T2 family. Red dot: All the T2 lines from this family carry the same mutation, indicating a mutation more likely inherited from the T1 parent rather than being *de novo* from the T2 line. Red square: Most active construct(s) for each panel.

In the combinations we tested, we detected low CRISPR activity using the meiosis I-specific promoter *MGE1* [26] (Fig 3C), the homeotic gene promoter *AG* [27] (Fig 3D) and the DNA polymerase subunit-encoding gene promoter *ICU2* [28] (Fig 3D). They were tested with constructs inducing an overall low activity and we do not exclude that they can perform efficiently in other conditions. In one context specifically, *ICU2* promoter resulted in moderate activity in four of the six T2 families tested, while only one T2 family showed activity with the *UBI10* promoter (Fig 3E).

*EC1.2* and an *EC1.2::EC1.1* fusion (referred as ‘*EC enhanced*’ or ‘*ECenh*’) are specifically expressed in the egg cell and were reported to trigger elevated mutation rates with CRISPR in Arabidopsis [17]. In our Golden Gate compatible system, only *ECenh* induced homozygous mutants in T1 and at low frequency (Fig 3B and G). In one comparison, *EC1.2* and *ECenh* performed slightly better than *pUBI10* (Fig 3D), but in another, they induced lower activity (Fig 3E).

A promoter from Cassava Vein Mosaic Virus (*pCsVMV*) was reported to mediate CRISPR activity in *Brassica oleracea* [29]. We found that it induced more CRISPR activity than *pUBI10* in two combinations tested (Fig 3D and E).

We also tested the *YAO* and *RPS5a* promoters. Both of them were reported to boost CRISPR activity in Arabidopsis [15,16]. Both triggered elevated mutation rates compared with the *UBI10* promoter (Fig 3F). In one comparison, *pRPS5a* performed slightly better (Fig 3G), but in another, *pYAO* performed better (Fig 3H).

As have others, we conclude that the promoter driving *Cas9* expression influences CRISPR-mediated mutation rates [15–17,26]. We observed the best mutation rates using *RPS5a, YAO* and *UBI10* promoters.

### Codon optimization of *Cas9* and presence of an intron elevate mutation rates

The activity of different constructs with the same promoter can be very different. For instance, *pRPS5a:Ca9* and *pYAO:Ca9* lines were recovered that displayed either high or low activity (Fig 3F and H). The most active constructs carried *Cas9_3* or *Cas9_4* alleles. We thus compared four *Cas9* alleles side-by-side (Fig 4). *Cas9_1* is a human codon-optimized version with a single C-terminal Nuclear Localization Signal (NLS) [3]. *Cas9_2* is an Arabidopsis codon-optimized version with a single C-terminal NLS [13]. *Cas9_3* is a plant codon-optimized version with both N- and C-terminal NLSs, an N-terminal FLAG tag and a potato intron *IV* [25]. *Cas9_4* is a human codon-optimized version with both N- and C-terminal NLSs and an N-terminal FLAG tag [10].

**Figure 4:**
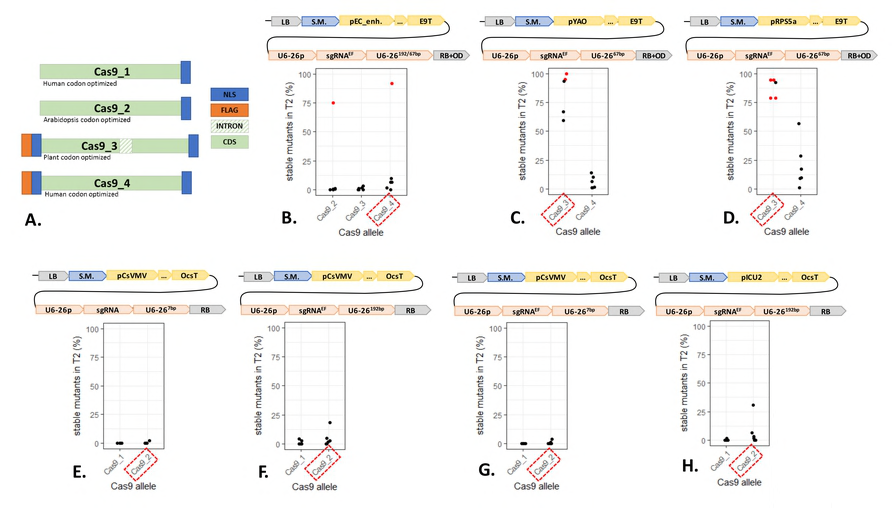
An intron-containing allele of *Cas9* triggers elevated mutation rates. **A.** <u>Cas9_1</u>: Mali *et al.*, 2013 [3]. <u>Cas9_2</u>: Fauser *et al.*, 2014 [13]. <u>Cas9_3</u>: Li *et al.*, 2013 [25]. <u>Cas9_4</u>: Le Cong *et al.*, 2013 [10]. <u>NLS</u>: Nuclear Localization Signal. <u>FLAG</u>: DYKDDDDK peptide. Apart from the FLAG and NLS, the amino acid sequences are identical. The nucleotide sequence (codon optimization) are different. Bars are not in scale. **B. to H.** Each panel represents a CDS comparison in the same context. CDSs can be compared within each panel, not from one panel to another. The modules were assembled into pICSL4723 (RB+OD, with an *overdrive*) or pAGM4723 (RB, without an *overdrive*) and transformed into Col-0 via *Agrobacterium tumefaciens* strain GV3101. <u>LB</u>: Left Border. <u>S.M</u>.: Selectable Marker (*Glufosinate resistance* gene). <u>pEC_enh.</u>: 752 bp of the At2g21740 promoter fused to 548 bp of the At1g76750 promoter. <u>pYAO</u>: 596 bp of the At4g05410 promoter. <u>pRPS5a</u>: 1688 bp of the At3g11940 promoter. <u>pCsVMV</u>: 517 bp of a promoter from *Cassava Vein Mosaic* virus. <u>pICU2</u>: 625 bp of the At5g67100 promoter. <u>E9T</u>: 631 bp of the *Pisum sativum rbcS E9* terminator. <u>OcsT</u>: 714 bp of the *Agrobacterium tumefaciens octopine synthase* terminator. <u>U6-26p</u>: 205 bp of the At3g13855 promoter. sgRNA^EF^: “extension-flip” sgRNA. <u>U6-26t</u>: 7, 67 or 192 bp of the At3g13855 terminator. <u>RB</u>: Right Border. For the comparison using the EC_enh. promoter, Cas9_2 is in combination with U6-26^192bp^; Cas9_3 and Cas9_4 are in combination with U6-26^67bp^. The sgRNA targets ADH1. CRISPR activity measured in % of homozygous or biallelic stable mutants in the second generation after transformation (T2). Each dot represents an independent T2 family. Red dot: All the T2 lines from this family carry the same mutation, indicating a mutation more likely inherited from the T1 parent rather than being *de novo* from the T2 line. Red square: Most active construct(s) for each panel.

We found that in comparable constructs, *Cas9_2* performs better than *Cas9_1* (Fig 4E to H), consistent with the fact that *Cas9_2* was designed for Arabidopsis codon usage. However, human codon-optimized *Cas9_4* induced more mutants than Arabidopsis optimized *Cas9_2* in one experiment (Fig 4B). Cas9_4 has an extra N-terminal NLS compared to Cas9_2, which may explain this difference. In this comparison specifically, *Cas9_3* was less efficient than *Cas9_4*. However, by comparing *Cas9_3* and *Cas9_4* in combination with *YAO* or *RPS5a* promoters, we found that *Cas9_3* resulted in high mutation rates (Fig 4C and D). *Cas9_3* efficiency can be explained by the plant codon optimization, the presence of two NLSs and the inclusion of a plant intron. This intron was originally added to avoid expression in bacteria during cloning and, as side effect, can also increase expression *in planta* [30]. We recommend the use of *Cas9_3* for gene editing in Arabidopsis.

### A modified sgRNA triggers CRISPR-induced mutations more efficiently

In the endogenous CRISPR immune system, Cas9 binds a CRISPR RNA (crRNA) and a trans-acting CRISPR RNA (tracrRNA) [31]. A fusion of both, called sgRNA, is sufficient for CRISPR-mediated genome editing [32]. sgRNA stability was suggested to be a limiting factor in CRISPR system [33]. Chen *et al.* proposed an improved sgRNA to tackle this issue [8]. It carries an A-T transversion to remove a TTTT potential termination signal, and an extended Cas9-binding hairpin structure (Fig 5A). We compared side-by-side the ‘<u>E</u>xtended’ and ‘<u>F</u>lipped’ sgRNA (sgRNA^EF^) with the classic sgRNA (Fig 5B and C). In two independent comparisons, the efficiency was higher with sgRNA^EF^. The improvement was not dramatic but sufficient to lead us to recommend use of ‘EF’-modified guide RNAs for genome editing in Arabidopsis.

**Figure 5:**
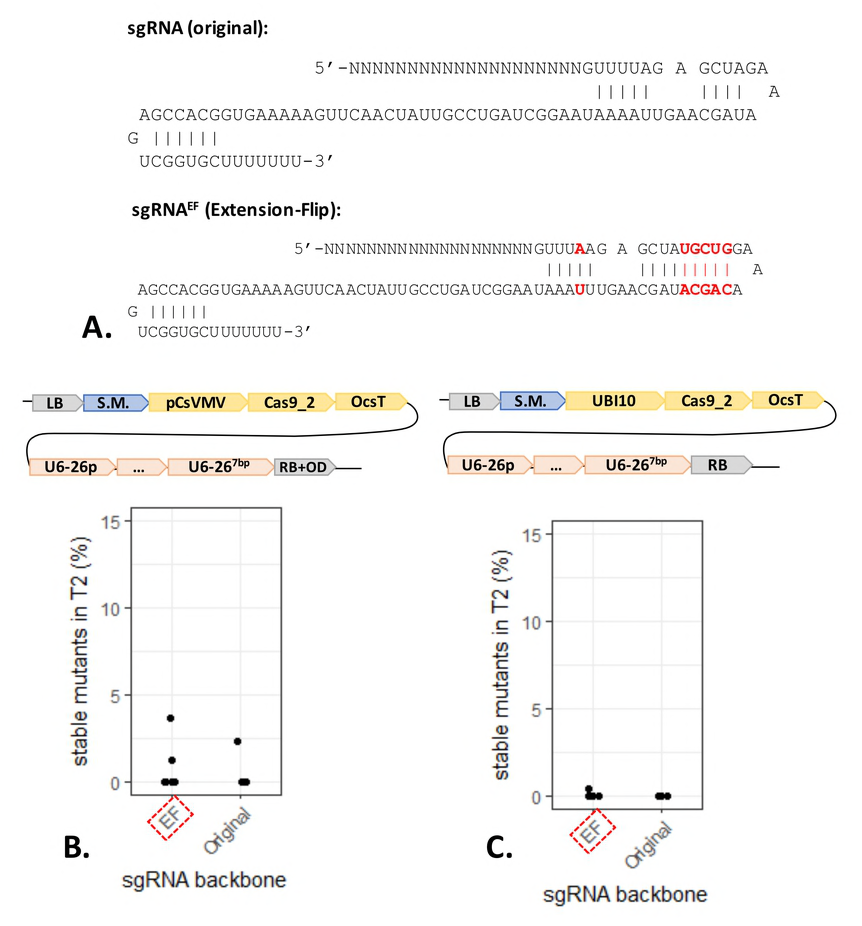
A modified sgRNA is slightly more efficient to trigger mutations. **A.** Original sgRNA proposed by Mali *et al*., 2013 [3]. Extension-Flip (EF) sgRNA proposed by Chen *et al*., 2013 [8]. **B. and C.** Each panel represents a sgRNA backbone comparison in the same context. sgRNA backbones can be compared within each panel but not from one panel to another. The modules were assembled into pICSL4723 (RB+OD, with an *overdrive*) or pAGM4723 (RB, without an *overdrive*) and transformed into Col-0 via *Agrobacterium tumefaciens* strain GV3101. <u>LB</u>: Left Border. <u>S.M</u>.: Selectable Marker (*Glufosinate resistance* gene). <u>pCsVMV</u>: 517 bp of a promoter from *Cassava Vein Mosaic* virus. <u>UBI10</u>: 1327 bp of the At4g05320 promoter. <u>Cas9_2</u>: Fauser *et al.*, 2014 [13]. <u>OcsT</u>: 714 bp of the *Agrobacterium tumefaciens octopine synthase* terminator. <u>U6-26p</u>: 205 bp of the At3g13855 promoter. <u>U6-26t</u>: 7 bp of the At3g13855 terminator. <u>RB</u>: Right Border. The sgRNA targets ADH1. CRISPR activity measured in % of homozygous or biallelic stable mutants in the second generation after transformation (T2). Each dot represents an independent T2 family. Red square: Most active construct(s) for each panel.

### The 3’ regulatory sequences of Cas9 and the sgRNA influence the overall activity

To avoid post-transcriptional modifications such as capping and polyadenylation, sgRNA must be transcribed by RNA polymerase III (Pol III). Several approaches involving ribozymes, Csy4 ribonuclease or tRNA-processing systems have been proposed but were not tested here [34–36]. *U6-26* is a Pol III-transcribed gene in Arabidopsis [37]. We used 205 bp of the 5’ upstream region of *U6-26* as promoter and we compared a synthetic polyT sequence (seven thymidines) and 192 bp of the 3’ downstream region as terminator. A T-rich stretch has been reported to function as a termination signal for Pol III [38].

In seven out of nine side-by-side comparisons, the authentic 192 bp of *U6-26* terminator directed a higher efficiency of the construct, as compared to a synthetic polyT termination sequence (Fig 6 and Fig S2). We speculate that a stronger terminator increases the stability of the sgRNA. For multiplex genome editing, the use of 192 bp per sgRNA will result in longer T-DNAs and increase the risk of recombination and instability. We generated constructs with only 67 bp of the *U6-26* 3’ downstream sequence. Such constructs were not compared side-by-side with the ‘192 bp terminator’, although they enabled high mutation rates (*e.g.* Fig 3F-G). Thus, we recommend using 67 bp of the 3’ downstream sequence of U6-26 as terminator for the sgRNA.

**Figure 6:**
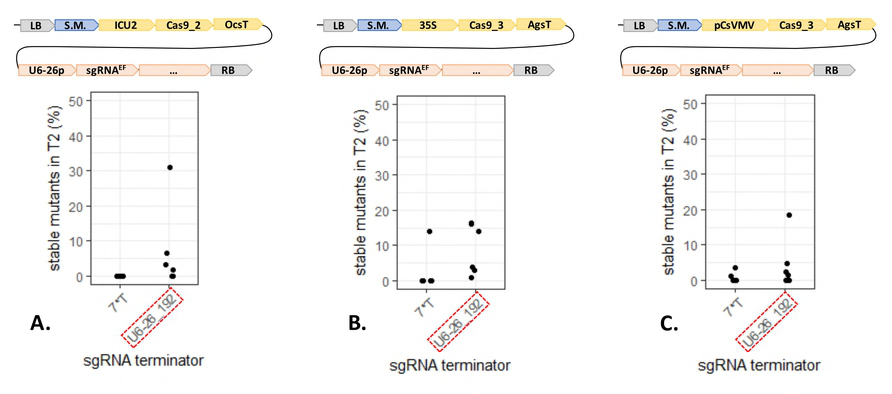
The sgRNA expression regulated by an authentic 3’ regulatory sequence of U6-26 produces greater mutation rates. **A. to C.** Each panel represents a terminator comparison in the same context. Terminators can be compared within each panel, not from one panel to another. The modules were assembled into pAGM4723 and transformed into Col-0 via *Agrobacterium tumefaciens* strain GV3101. <u>LB</u>: Left Border. <u>S.M</u>.: Selectable Marker (*Glufosinate resistance* gene). <u>ICU2</u>: 625 bp of the At5g67100 promoter. <u>35S</u>: 426 bp of the 35S promoter from Cauliflower Mosaic Virus. <u>pCsVMV</u>: 517 bp of a promoter from *Cassava Vein Mosaic* virus. <u>Cas9_2</u>: Fauser *et al.*, 2014 [13].<u>Cas9_3</u>: Li *et al.*, 2013 [25]. <u>OcsT</u>: 714 bp of the *Agrobacterium tumefaciens octopine synthase* terminator. <u>AgsT</u>:410 bp of the *Agrobacterium tumefaciens agropine synthase* terminator. <u>U6-26p</u>: 205 bp of the At3g13855 promoter. sgRNA^EF^: “extension-flip” sgRNA. <u>U6-26t</u>: 7 or 192 bp of the At3g13855 terminator. <u>RB</u>: Right Border. The sgRNA targets ADH1. CRISPR activity measured in % of homozygous or biallelic stable mutants in the second generation after transformation (T2). Each dot represents an independent T2 family. Red square: Most active construct(s) for each panel.

Since 3’ regulatory sequences can influence sgRNA stability, we tested if the same was true for *Cas9*. We compared the *Pisum sativum rbcS E9* with two *A. tumefaciens* terminators commonly used in Arabidopsis: *Ocs* and *Ags* (Fig 7). We did not observe consistent differences between *E9* and *Ocs* (Fig 7A and B). However, in one comparison, *E9* outperformed *Ags* (Fig 7C). This is consistent with previous observations that RNA Polymerase II (Pol II) terminators quantitatively control gene expression and influence CRISPR efficiency in Arabidopsis [17,39]. We propose that a weak terminator after *Cas9* enables Pol II readthrough that could interfere with Pol III transcription of sgRNAs in some T-DNA construct architectures. This limiting factor can be corrected by divergent transcription of *Cas9* and *sgRNAs*.

**Figure 7:**
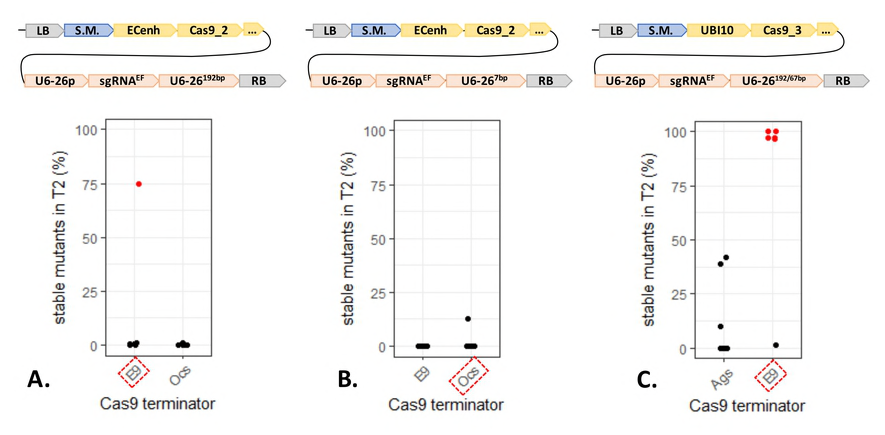
A weak 3’ regulatory sequence reduces the CRISPR-induced mutation rate. **A. to C.** Each panel represents a terminator comparison in the same context. Terminators can be compared within each panel, not from one panel to another. The modules were assembled into pAGM4723 and transformed into Col-0 via *Agrobacterium tumefaciens* strain GV3101. <u>LB</u>: Left Border. <u>S.M</u>.: Selectable Marker (*Glufosinate resistance* gene). <u>EC_enh.</u>: 752 bp of the At2g21740 promoter fused to 548 bp of the At1g76750 promoter. <u>UBI10</u>: 1327 bp of the At4g05320 promoter. <u>Cas9_2</u>: Fauser *et al.*, 2014 [13]. <u>Cas9_3</u>: Li *et al.*, 2013 [25]. <u>E9T</u>:631 bp of the *Pisum sativum rbcS E9* terminator. <u>OcsT</u>: 714 bp of the *Agrobacterium tumefaciens octopine synthase* terminator. <u>AgsT</u>: 410 bp of the *Agrobacterium tumefaciens agropine synthase* terminator. <u>U6-26p</u>: 205 bp of the At3g13855 promoter. sgRNA^EF^: “extension-flip” sgRNA. <u>U6-26t</u>: 7, 67 or 192 bp of the At3g13855 terminator. <u>RB</u>: Right Border. For the comparison using the UBI10 promoter, the AgsT is in combination with U6-26^192bp^; OcsT is in combination with U6-26^67bp^. The sgRNA targets ADH1. CRISPR activity measured in % of homozygous or biallelic stable mutants in the second generation after transformation (T2). Each dot represents an independent T2 family. Red dot: All the T2 lines from this family carry the same mutation, indicating a mutation more likely inherited from the T1 parent rather than being *de novo* from the T2 line. Red square: Most active construct(s) for each panel.

### Divergent transcription of Cas9 and sgRNA expression can elevate mutation rates

The Golden Gate Level 1 acceptor vector collection contains seven ‘forward’ expression cassettes and seven ‘reverse’ expression cassettes, which are interchangeable [23]. We assembled ‘*RPS5a:Cas9_4:E9*’ and ‘*YAO:Cas9_3:E9*’ in both the Level 1 vector position 2 forward (pICH47742) and reverse (pICH47811) (Fig 1 and 6). In one case, CRISPR activity was moderate when *Cas9* and *sgRNA* are expressed in the same direction and high when they are expressed in opposite direction (Fig 8A). In another case, CRISPR activity was very high in both cases (Fig 8B).

**Figure 8:**
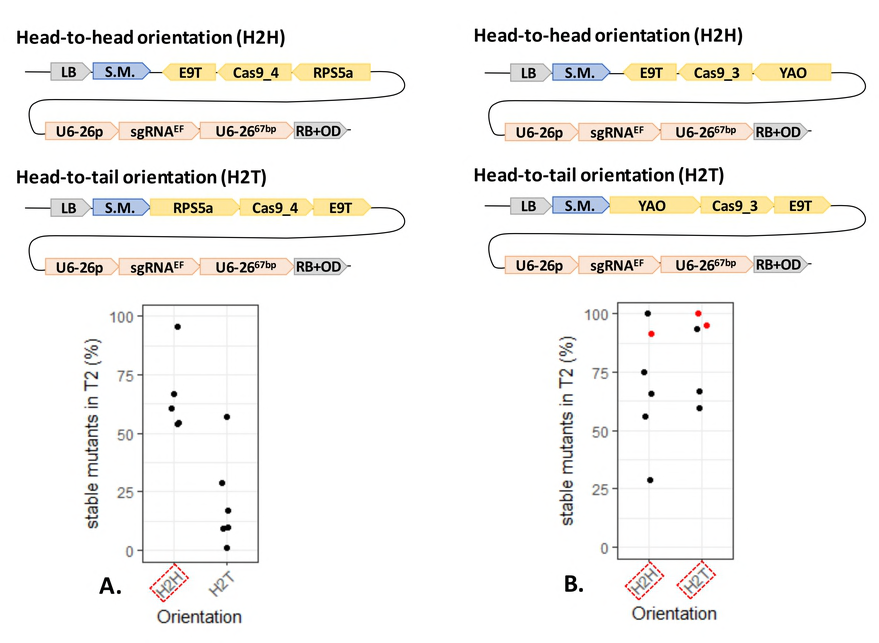
CRISPR activity is similar or higher when the sgRNA and the Cas9 expression cassettes are in a head-to-head orientation. **A. and B.** Each panel represents an orientation comparison in the same context. Orientations can be compared within each panel, not from one panel to another. The modules have been assembled by Golden Gate into pICSL4723 (RB+OD, with an *overdrive*) and transformed into Col-0 via *Agrobacterium tumefaciens* strain GV3101. <u>LB</u>: Left Border. <u>S.M</u>.: Selectable Marker (*Glufosinate resistance* gene. <u>RPS5a</u>: 1688 bp of the At3g11940 promoter. <u>YAO</u>: 596 bp of the At4g05410 promoter. <u>Cas9_3</u>: Li *et al.*, 2013 [25]. <u>Cas9_4</u>: Le Cong *et al.*, 2013 [10]. <u>E9T</u>: 631 bp of the *Pisum sativum rbcS E9* terminator. <u>U6-26p</u>: 205 bp of the At3g13855 promoter. sgRNA^EF^: “extension-flip” sgRNA. <u>U6-26t</u>: 67 bp of the At3g13855 terminator. <u>RB</u>: Right Border. The sgRNA targets ADH1. CRISPR activity measured in % of homozygous or biallelic stable mutants in the second generation after transformation (T2). Each dot represents an independent T2 family. Red dot: All the T2 lines from this family carry the same mutation, indicating a mutation more likely inherited from the T1 parent rather than being *de novo* from the T2 line. Red square: Most active construct(s) for each panel.

We thus recommend to both use a strong terminator after *Cas9* (*e.g. E9* or *Ocs*) and express *Cas9* and *sgRNA* in opposite directions.

## DISCUSSION

CRISPR emerged in 2012 as a useful tool for targeted mutagenesis in many organisms including plants [11,32]. In Arabidopsis, the transgenic expression of CRISPR components can be straightforward, avoiding tedious tissue culture steps. Many strategies to enhance the overall CRISPR-induced mutation rate have been proposed [8,13,15–17,40]. Here we report a systematic comparison of putative limiting factors including promoters, terminators, codon optimization, sgRNA improvement and T-DNA architecture.

We found that the best promoters to control Cas9 expression are *UBI10, YAO* and *RPS5a*. The best terminators in our hands were *Ocs* from *A. tumefaciens* and *rbcS E9* from *P. sativum.* A plant codon-optimized, intron-containing *Cas9* allele outperformed the other alleles tested. A modified sgRNA with a hairpin Extension and a nucleotide Flip, called sgRNA^EF^, triggers slightly elevated mutations rates. The sgRNA transcription regulation by the authentic 3’ regulatory sequence of *AtU6-26* results in better CRISPR activity. We get high mutation rates with either 67 bp or 192 bp of terminator and recommend using the shortest (67 bp). We hypothesise that a weak terminator after *Cas9* enables RNA-polymerase II readthrough within the sgRNA expression cassette, preventing optimal expression of the sgRNA. We indeed elevate CRISPR efficiency by expressing *Cas9* and *sgRNA* in opposite directions.

Finally, we recommend to use a ‘*YAO:Cas9_3:E9*’ and a ‘*pU6-26:sgRNA*^*EF*^:*U6-26t*^*67bp*^’ cassettes in head-to-head orientation. This combination is included in the constructs tested here (Fig 8B) and enabled us to recover two homozygous mutants in five T1 plants tested. We also obtained useful rates with other constructs (*e.g.* Fig 3F), indicating that the CRISPR components do not entirely explain the final CRISPR activity. It was recently reported that heat stress increases the efficiency of CRISPR in Arabidopsis [41]. Plants grown at different times (*e.g.* winter or summer) might experience slightly different environments. These differences may explain fluctuation of the CRISPR activity that we observed, independently of the T-DNA architecture.

From the mutant screen, 315 allyl-alcohol resistance lines were confirmed by capillary sequencing (Table S5). We classified them in four categories: (i) 59% were homozygous (single sequencing signal, different than *ADH1* WT), (ii) 11% were heterozygous (dual sequencing signal, one matching *ADH1* WT), (iii) 10% were biallelic (dual sequencing signal, none matching *ADH1* WT) and (iv) 20% were difficult to assign (unclear sequencing signals, either biallelic or due to somatic mutations, but clearly different than WT, heterozygous or homozygous genotypes) (Fig 9). The recovery of heterozygous (*ADH1/adh1*) lines indicates that the loss of a single copy of *ADH1* can sometimes enable plants to survive the allyl-alcohol selection.

**Figure 9:**
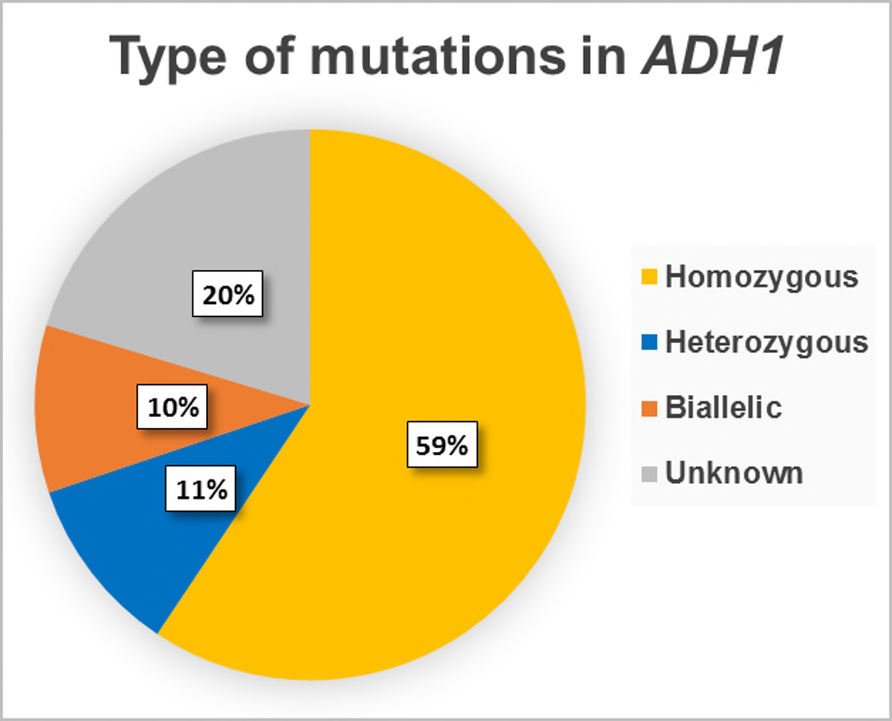
Genotype at *ADH1* locus confirmed by capillary sequencing. For each T2 family tested, up to six allyl-alcohol resistant plants were genotyped by capillary sequencing of an sgRNA target (*ADH1*) PCR amplicon. We retrieved a total of 315 sequences with a mutation. 59% (187) showed a single sequencing signal, different than *ADH1* WT and were classified as “Homozygous”. 11% (33) showed an overlap of two sequencing signals, one matching *ADH1* WT and one different; and were classified as “Heterozygous”. 10% (31) showed an overlap of two sequencing signals, none matching *ADH1* WT; and were classified as “Biallelic”. 64 (20%) showed an overlap of signals different than WT but not clear enough to distinguish; and were classified as “Unknown”. The “Unknown” sequences can be biallelic or due to somatic mutations but are different than WT, heterozygous or homozygous genotypes. The figure was made using Excel (Microsoft Office 365 ProPlus).

Stable double mutations are the result of two CRISPR events, on the male and female inherited chromosome respectively. In this scenario, lines can be recovered with two different mutations, resulting in a biallelic (*e.g. adh1-2/adh1-3*) genotype, rather than having the same mutation on both chromosomes (*e.g. adh1-1/adh1-1*). Surprisingly, we recovered more homozygous than biallelic events. The simplest explanation for an excess of homozygotes is that after CRISPR-induced mutation of one *ADH1* copy, Cas9 cleavage of the remaining WT copy of *ADH1* is processed by somatic recombination resulting in gene conversion from the first mutation. Indeed, intrachromosomal somatic recombination occurs in plants and is enhanced by ionizing radiation which can cause double strand breaks [42]. The prevalence of homozygous over biallelic genotypes facilitates the genotyping and is an advantage for targeted mutagenesis using CRISPR-Cas9.

We used a glufosinate resistance selectable marker which enables easy selection of transgenic lines. It can be important to segregate away the T-DNA in the CRISPR mutant line for multiple reasons. For instance, a loss-of-function phenotype must be confirmed by complementation of the CRISPR-induced mutation. A CRISPR construct still present in the mutant can target the complementation transgene and interfere with the resulting phenotypes. Selection of non-transgenic lines is possible but complicated with classic selectable markers such as kanamycin or glufosinate resistance, since a selective treatment kills the non-transgenic plants. FAST-Green and FAST-Red provide a rapid non-destructive selectable marker and involve expression of a GFP-or RFP-tagged protein in the seed [43]. Transgenic and non-transgenic seeds can be distinguished under fluorescence microscopy. This facilitates recovery of mutant seed lacking the T-DNA (Fig 10). Homozygous mutants can be identified among the independent T1 lines. Non-fluorescent seeds can be selected from the T1 seeds. The resulting T2 plants are homozygous mutant and non-transgenic.

**Figure 10:**
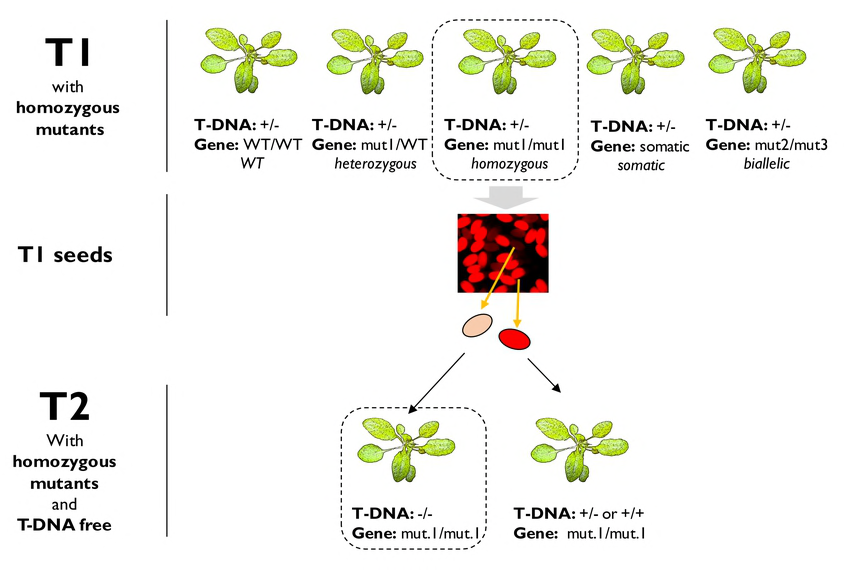
FAST-Red combined with CRISPR to generate T-DNA free mutants. The six T1 lines are independent transformants. They are all hemizygous for the T-DNA. At the sgRNA target site, they can be WT, or display somatic, heterozygous, biallelic of homozygous mutations. All the possibilities are represented here. One line has homozygous mutation (*mut1/mut1*). It produces seeds segregating for the T-DNA, visible under microscope if using FAST-Red. The seeds will segregate 3:1 (Red : Non-red) if there is one locus insertion, 15:1 (Red : Non-red) if there are two loci insertion, etc. The T2 progeny of (mut1/mut1) is 100% homozygous for the mutation. The non-red seeds are also T-DNA free.

We report here a CRISPR- and Golden Gate-based method to generate stable Arabidopsis mutant lines in one generation. In our efforts to elevate mutation rates in Arabidopsis, we found several limiting factors mostly related to *Cas9* and *sgRNA* transcription. Some of these findings can be tested for other plant species and for knock-in breeding. The generation of null alleles via CRISPR is today quick and simple, facilitating the investigation of gene function. Improvement of rates of gene ‘knock-ins’ provides the next challenge. *In vivo* gene tagging or knock-in breeding are theoretically possible and have been reported [44,45]. Improvements in CRISPR-based genome editing techniques will facilitate the study of genes and proteins and be beneficial for both basic and applied plant science

## MATERIALS AND METHODS

### CRISPR constructs assembly

The vectors were assembled using the Golden Gate modular cloning method [23]. To generate the Cas9 expression cassettes, the *RPS5a, YAO, ICU2, CsVMV, EC, EC_enh*., *UBI10, AG, MGE1* and *35S* promoters, the *Cas9_1, Cas9_2, Cas9_3* and *Cas9_4* coding sequences, the *Ocs, Nos, Ags* and *E9* terminators were amplified using primers flanked with BpiI restriction sites associated with Golden Gate compatible overhangs (Table S3). 0.02 pmoles of the purified PCR products were mixed with the same molar amount of the corresponding Level 0 vector (Table S3), 0.5 µl of BpiI enzyme (10U/µl, ThermoFisher), 0.5 µl of T4 ligase (400U/µl, NEB), 1.5 µl of CutSmart Buffer (NEB), 1.5µl of Bovine Serum Albumin (10X) and water in a total reaction volume of 15 µl. The reaction was placed in a thermocycler and the following ‘Golden Gate’ program was applied: 20 seconds 37°C, 25 cycles of [3 minutes 37°C / 4 minutes 16°C], 5 minutes 50°C and 5 minutes 80°C.

Combinations of three Level 0 vectors containing respectively a promoter, a Cas9 coding sequence and a terminator were assembled in Level 1 vector pICH7742 (Position 2) or pICH47811 (Position 2, reverse) by the same ‘Golden Gate’ protocol but using 0.5 µl of BpiI enzyme (10U/µl, ThermoFisher) instead of 0.5 µl of BsaI-HF.

To generate the sgRNA expression cassettes, DNA fragments containing the classic or the ‘EF’ backbone with 7, 67 or 192 bp of the *U6-26* terminator were amplified using primers flanked with BsaI restriction sites associated with Golden Gate compatible overhangs (Table S3). The amplicons were assembled with the U6-26 promoter (pICSL90002) in Level 1 vector pICH7751 (Position 3) by the ‘Golden Gate’ protocol using the BsaI-HF enzyme. Combinations of three Level 1 vectors containing a glufosinate resistance selectable maker (pICSL11017), a Cas9 expression cassette and a sgRNA expression cassette were assembled in Level 2 pAGM4723 (without an *overdrive*) or pICSL4723 (with an *overdrive*) by the ‘Golden Gate’ protocol using the BpiI enzyme. All the plasmids were prepared using a QIAPREP SPIN MINIPREP KIT on *Escherichia coli* DH10B electrocompetent cells selected with appropriate antibiotics and X-gal.

All the plasmid identification numbers refer to the ‘addgene database’ (www.addgene.org/).

### Plant transformation, growth and selection

*Agrobacterium tumefaciens* strain GV3101 was transformed with plasmids by electroporation and used for stable transformation of Arabidopsis accession Col-0. Arabidopsis plants were grown in ‘short days’ conditions (10 hr light/14 hr dark, 21°C). Transformants were selected by spraying three times 1- to 3-weeks old seedlings with phosphinotrycin at a concentration of 0.375g/l. 4-weeks old resistant plants were transferred in ‘long days’ conditions (16 hr light/8 hr dark, 21°C) for flowering. For each genotype, six independent T1 were self-pollinated to obtain six independent T2 families per construct.

### Characterisation of CRISPR events

T2 families were tested for resistance to allyl-alcohol. ∼100 seeds were sterilized, immersed in water (4°C, dark, overnight), treated with allyl-alcohol (30mM, room temperature, 2 hours, shaken at 750rpm), rinsed three times with water and sown on MS^1/2^ medium. After two weeks, the number of germinated and non-germinated seeds was monitored. DNA was extracted from up six allyl-alcohol resistant plants (or all the resistant plants if there were less than six) for genotyping. ∼0.5cm^2^ of leaf tissue was stuck FTA filter paper (Whatman^TM^ Bioscience). 1-mm disks were punched out from FTA filter paper by using a punch and placed in a 200µl PCR tubes. Samples were incubated in 50µl of FTA buffer (1.25ml Tris 1M, 500µl EDTA 0.5M, 12.5µl Tween 20 and water up to a total volume of 125ml) for 2 hours and rinsed with water. PCR was performed on this template using primers flanking the sgRNA target in *ADH1* (Table S3) and Q5^®^ High-Fidelity DNA Polymerase (NEB, following the manufacturer recommendations). After amplification, the PCR products were resolved by electrophoresis on a 1.5% agarose gel and purified using the QIAquick Gel Extraction Kit (QIAGEN). The purified PCR product was sequenced using the same primer set for amplifications by capillary sequencing (GATC Biotech). Sequencing results were compared to the Col-0 sequence of *ADH1* using CLC Main Workbench 7.7.1. *ADH1* genotypes were reported as WT (identical to Col-0), heterozygous (both Col-0 and single mutation detected), biallelic (two different mutations detected), homozygous (single mutation detected) or somatic (more than two signals detected). The number of confirmed mutant among all the allyl-alcohol resistant lines was used to estimate the total number of real mutants among allyl-alcohol survivors from each plate. For each T2 family, the CRISPR efficiency was defined as the ratio of homozygous and biallelic mutants compared to the total number of seeds sown. Plots presented in this article were made using *ggplot2* in *R* version 3.3.2.

## ACKNOWLEDGEMENT

This work was supported by the Gatsby Charitable Foundation at The Sainsbury Laboratory, Norwich, and the Bill and Melinda Gates Foundation (Grand Challenges Exploration, grant agreement OPP1060026). Federica LOCCI was supported by the “Funding of joint research projects for the PhD students’ mobility abroad” from Sapienza University of Rome. The funders had no role in study design, data collection and analysis, decision to publish, or preparation of the manuscript. We thank Dr Sylvestre Marillonnet for help refining Golden Gate cloning vectors.

## AUTHOR CONTRIBUTIONS

Conceptualization: BC, LT, YY and JDGJ. Data curation: BC.

Formal analysis: BC.

Funding acquisition: JDGJ.

Investigation: LT, FL and BC.

Methodology: BC, LT, YY and JDGJ.

Project administration: JDGJ.

Resources: LT, YY, FL and BC.

Supervision: JDGJ.

Visualization: BC.

Writing – original draft: BC.

Writing – review & editing: BC, LT and JDGJ.

## SUPPORTING INFORMATION

**Table S1:** List of the Level 2 T-DNA used in this article.

**Table S2:** List of plasmids used in this article, available through addgene.

**Table S3:** List of primers used in this article. Some vectors were not cloned using a PCR step (*e.g.* synthesised or cloned prior this article), which are indicated in this table.

**Table S4:** Mutation rates table. LBC number indicates a unique independent T1 line. Clone, SLJ number and Genotype refers to the “Level 2 Constructs” table (Table S1). Vector “pAGM4723” lack an *overdrive*; Vector “pICH4723” has an *overdrive*. “Same_mutation” indicates whether all the lines carry the same mutation. It is applied only if more than 75% of the seeds germinated. If so, it indicates that the parent was likely a homozygous mutant and the mutation was inherited to all progenies.

**Table S5:** List of mutations obtained in *ADH1* (from capillary sequencing data).

**Figure S1. The presence of the *overdrive* sequence in the T-DNA Right Border does not affect the CRISPR efficiency**

**A.** Sequence of the right border with (pICSL4723) or without (pAGM4723) the *overdrive* sequence. **B. and C.** Each panel represents a vector comparison in the same context. Vectors can be compared within each panel, not from one panel to another. The modules have been assembled by Golden Gate into pICSL4723 (OD+, with an *overdrive*) or pAGM4723 (OD-, without an *overdrive*) and transformed into Col-0 via *Agrobacterium tumefaciens* strain GV3101. <u>LB</u>: Left Border. <u>S.M</u>.: Sel. Marker (Glufosinate *resistance* gene). <u>EC</u>: 1014 bp of the At2g21740 promoter. <u>EC_enh.</u>: 752 bp of the At2g21740 promoter fused to 548 bp of the At1g76750 promoter. <u>Cas9_2</u>: Fauser *et al.*, 2014 [13]. <u>E9T</u>: 631 bp of the *Pisum sativum rbcS E9* terminator. <u>U6-26p</u>: 205 bp of the At3g13855 promoter. sgRNA^EF^: “extension-flip” sgRNA. <u>U6-26t</u>: 7 bp of the At3g13855 terminator. <u>RB</u>: Right Border. The sgRNA targets ADH1. CRISPR activity measured in % of homozygous or biallelic stable mutants in the second generation after transformation (T2). Each dot represents an independent T2 family. Red square: Most active construct(s) for each panel. The *overdrive* sequence can increase the integration efficiency [21]. In one comparison the presence of the *overdrive* results in slightly better activity (C), but in another one it did not (B). We concluded that the presence of an *overdrive* does not influence the CRISPR efficiency. Thus, we could compare constructs independently of the presence of an *overdrive*.

**Figure S2: The sgRNA expression regulated by an authentic 3’ regulatory sequence of U6-26 produces greater mutation rates**

**A. To F.** Each panel represents a terminator comparison in the same context. Terminators can be compared within each panel, not from one panel to another. The modules were assembled into pAGM4723 and transformed into Col-0 via *Agrobacterium tumefaciens* strain GV3101. <u>LB</u>: Left Border. <u>RB</u>: Right Border. <u>S.M</u>.: Sel. Marker (Glufosinate *resistance* gene). <u>CsVMV</u>: 517 bp of a promoter from Cassava Vein Mosaic virus. <u>UBI10</u>: 1327 bp of the At4g05320 promoter. <u>EC</u>: 1014 bp of the At2g21740 promoter. <u>EC_enh.</u>: 752 bp of the At2g21740 promoter fused to 548 bp of the At1g76750 promoter. <u>Cas9_1</u>: Mali *et al.*, 2013 [3]. <u>Cas9_2</u>: Fauser *et al.*, 2014 [13]. <u>E9T</u>: 631 bp of the *Pisum sativum rbcS E9* terminator. <u>OcsT</u>: 714 bp of the *Agrobacterium tumefaciens octopine synthase* terminator. <u>EF</u>: 205 bp of the At3g13855 promoter controlling the expression of an “extension-flip” sgRNA. <u>U6-26t</u>: 7 or 192 bp of the At3g13855 terminator. The sgRNA targets ADH1. CRISPR activity measured in % of homozygous or biallelic stable mutants in the second generation after transformation (T2). Each dot represents an independent T2 family. Red dot: All the T2 lines from this family carry the same mutation, indicating a mutation more likely inherited from the T1 parent rather than being *de novo* from the T2 line. Red square: Most active construct(s) for each panel.

